# The antimicrobial peptide TAT-RasGAP_317-326_ inhibits the formation and the expansion of bacterial biofilms *in vitro*

**DOI:** 10.1101/2020.09.29.318378

**Authors:** Tytti Heinonen, Simone Hargraves, Maria Georgieva, Christian Widmann, Nicolas Jacquier

**Affiliations:** Department of Laboratories, Institute of Microbiology, Lausanne University Hospital and University of Lausanne, Lausanne, Switzerland; Department of Physiology, University of Lausanne, Lausanne, Switzerland

**Keywords:** biofilms, antimicrobial peptide, antibiotic resistance, TAT-RasGAP_317-326_

## Abstract

Biofilms are structured aggregates of bacteria embedded in a self-produced matrix. Pathogenic bacteria can form biofilms on surfaces and in tissues leading to nosocomial and chronic infections. While antibiotics are largely inefficient in limiting biofilm formation and expansion, antimicrobial peptides (AMPs) are emerging as alternative anti-biofilm treatments. In this study, we explore the effect of the newly described AMP TAT-RasGAP_317-326_ on *Acinetobacter baumannii, Pseudomonas aeruginosa* and *Staphylococcus aureus* biofilms. We observe that TAT-RasGAP_317-326_ inhibits the formation of biofilms at concentrations equivalent or two times superior to the minimal inhibitory concentration (MIC) of the corresponding planktonic bacteria. Moreover, TAT-RasGAP_317-326_ limits the expansion of *A. baumannii* and *P. aeruginosa* established biofilms at concentrations 2-4 times superior to the MIC. These results further confirm the potential of AMPs against biofilms, expand the antimicrobial potential of TAT-RasGAP_317-326_ and support further development of this peptide as an alternative antimicrobial treatment.

## Introduction

The emergence of antibiotic resistance is a major threat to public health. Infections caused by multidrug resistant bacteria are an increasing concern, leading to disabilities and possibly death [1]. One limitation in the development of novel antibiotics is the use of free-living (also called planktonic) bacteria in axenic medium as model system. This model is not representative of infections, as the majority of them are caused by multicellular aggregates of bacteria in a self-produced matrix, called biofilms. [2]. The formation of biofilms is dynamic and triggered by signals such as nutrient limitation, antibiotic exposure and oxygen availability [3]. The matrix embedding the biofilm is composed of polysaccharides, DNA and proteins, and forms a scaffold for bacterial attachment, protects bacteria from external insults, and enables a compartmentalisation of the biofilms with distinct bacterial subpopulations [2, 4].

Pathogenic bacteria, including *Acinetobacter baumannii, Pseudomonas aeruginosa* and *Staphylococcus aureus*, can form biofilms on medical devices implanted in humans, develop in some tissues, e.g. lungs, teeth and skin, and may persist on healthcare surface [5, 6]. Moreover, bacteria present in biofilms can be 10 to1000 times more resistant to antibiotics than planktonic bacteria [7]. Bacteria contained in biofilms strongly differ from planktonic bacteria regarding both their gene expression profile and their functional properties. Moreover, bacterial subpopulations composing biofilms are heterogeneous, further complicating biofilm treatment. Current biofilm treatment is tedious and resembles cancer therapy starting with the surgical removal of the biofilm followed by administration of antimicrobials [8]. This strategy is not optimal since antimicrobials are often unable to eradicate all bacterial subpopulations embedded in the biofilm. Moreover, this treatment may not completely disrupt the biofilm scaffold, potentially allowing its colonization by other microorganisms. The combination of intrinsic antibiotic resistance with resistance properties provided by the biofilm organisation renders these nosocomial infections highly challenging to treat. We thus need alternatives to classical antibiotics to treat biofilms formed by resistant pathogens.

Antimicrobial peptides (AMPs) could be such an alternative. AMPs are short peptides (up to 100 amino acids) produced by all living organisms as a defence mechanism that usually target a broad range of pathogens [9]. While the ability of bacteria to develop resistance towards AMPs is debated [10], pathogens carrying resistance genes to one or multiple antibiotics often show increased sensitivity towards AMPs [11, 12]. Some AMPs, alone or combined with antibiotics, act both on the inhibition of biofilm formation and on the disruption of mature biofilm [13].

TAT-RasGAP_317-326_ is a chimeric peptide consisting of the cell-permeable HIV peptide TAT_48-57_ linked to a 10-amino acid sequence of the Src Homology 3 Domain (SH3 domain) of p120 RasGAP [14]. TAT-RasGAP_317-326_ was first described for its anticancer properties, being able to sensitise cancer cells to anticancer therapies [15-17], directly kill cancer cells [18] and display anti-metastatic effect [19]. More recently, we showed that TAT-RasGAP_317-326_ exerts a broad antimicrobial activity against both Gram-positive and Gram-negative human pathogens including *A. baumannii, P. aeruginosa* and *S. aureus* [20]. Like other AMPs, TAT-RasGAP_317-326_ is positively charged and contains several arginine and tryptophan residues. Arginine residues may interact with the negatively charged bacterial membrane while tryptophan may insert in the bacterial membrane [21, 22]. These interactions may enable AMPs to translocate in the bacteria without disrupting membrane and to target intracellular components. While the tryptophan at position 317 of RasGAP domain is essential for the antimicrobial activity of TAT-RasGAP_317-326_, the mode of action of this peptide remains unknown.

In this report, we questioned the potential effect of TAT-RasGAP_317-326_ on biofilm formation and on mature biofilm expansion. We show that TAT-RasGAP_317-326_ inhibits the formation of *A. baumannii, P. aeruginosa* and *S. aureus* biofilms *in vitro* at concentrations equal or 2 times the MIC of planktonic bacteria. Moreover, treatment of mature biofilms with TAT-RasGAP_317-326_ reduces the expansion of *A. baumannii* and *P. aeruginosa* biofilms, but, similarly to other tested antibiotics and AMPs, cannot completely eradicate the biofilm scaffold. These results highlight the potential of TAT-RasGAP_317-326_ in the prevention and treatment of biofilms and encourage further development of AMPs as alternative antimicrobial compounds.

## Material and methods

### Bacterial strains and growth conditions

*Acinetobacter baumannii* ATCC 19606 (*A. baumannii*) and *Staphylococcus aureus* ATCC 29213 (*S. aureus*) were from American Type Culture Collection (ATCC; Manassas, VA). *Pseudomonas aeruginosa* PA14 (*P. aeruginosa*) was obtained from Professor Leo Eberl (Department of Plant and Microbial Biology, University of Zürich, Switzerland). Bacteria were routinely grown in Mueller-Hinton broth (*A. baumannii*), tryptic soy broth (TSB; *S. aureus*) and Luria-Bertani broth (LB; *P. aeruginosa*).

### Peptides and antibiotics

TAT-RasGAP_317-326_ is a retro-inverso peptide (i.e. reversed direction compared to natural sequence including D-amino acids) with an antimicrobial activity [20] composed of amino acids 48-57 of HIV TAT protein (RRRQRRKKRG) and 317-326 of human RasGAP protein (DTRLNTVWMW) linked with two glycines. TAT-RasGAP_317-326_ was synthesized by SBS Genetech (Beijing, China). Ciprofloxacin, tetracycline and gentamicin were from Applichem (Darmstadt, Germany), rifampicin and polymyxin B from Sigma-Aldrich (Saint-Louis, MO), and melittin from Enzo Life Sciences (Farmingdale, NY).

### MIC measurement

The minimal inhibitory concentrations (MICs) of antibiotics and AMPs on bacterial strains were determined as described [23]. Briefly, overnight cultures were diluted to 0.1 OD_600_, grown for 1h at 37°C with 200 rpm shaking and diluted 1:200 in 96-well plates containing increasing amount of antibiotics or AMPs. Plates were incubated statically for 18h at 37°C. The lowest concentration at which no turbidity could be observed was determined as MIC.

### Biofilm formation assay and treatment

Overnight cultures of bacteria were diluted 1:50 and grown to exponential phase at 37°C. Cultures were washed with PBS and adjusted to 0.1 OD_600_ (10^7^-10^9^ CFU/ml) in BM2 medium (62 mM potassium phosphate buffer, 7 mM ammonium sulfate, 10 µM iron sulfate, 0.4% (w/v) glucose, 0.5% (w/v) casaminoacids, 2 mM magnesium sulfate) with or without antibiotics or AMPs. Hundred microliters of culture were plated in polypropylene plates (Greiner, Kremsmünster, Austria) and biofilms were allowed to form for 24h (*A. baumannii* and *P. aeruginosa*) and 48h (*S. aureus*) at room temperature (RT). For biofilm eradication, mature biofilms were washed twice with PBS and incubate with antibiotics or AMPs in BM2 medium for 24h at RT.

### Measurement of biofilm biomass and bacterial viability

To assess total biofilm biomass, biofilms were washed with water and stained with 0.1% crystal violet (Sigma-Aldrich) as described [24]. Stained biofilms were dried overnight at RT and dissolved in 30% acetic acid. Absorbance was measured at 590 nm. To assess bacterial viability, biofilms were washed twice with PBS and incubated with 4 µg/ml resazurin (Sigma-Aldrich) in BM2. Plates were incubated 90 minutes at 37°C. Fluorescence was measured with a FLUOstar Omega plate reader (BMG Labtech, Ortenberg, Germany) with Ex/Em = 540/580 and gain = 500.

### Calculations

The antibiotic or AMP concentration added at biofilm induction that reduced the formation of biofilm (viability and biomass) by ≥ 90% was set as BPC_90_ (biofilm prevention concentration). For mature biofilms, the MBIC (minimal biofilm inhibitory concentration) was set as the concentration of antibiotic or AMP that resulted in no expansion of an existing biofilm (viability and biomass equal or lower than the value at treatment). The MBEC_90_ (minimal biofilm eradication concentration) was defined as the concentration of antibiotic or AMP that reduced the biofilm initial biomass by ≥ 90% during treatment. All calculations were adapted from [25].

## Results

### Classical antibiotics have a moderate effect on *A. baumannii* and *P. aeruginosa* biofilms

*A. baumannii* and *P. aeruginosa* form biofilms that show increased resistance to antibiotics compared to planktonic bacteria [25, 26]. We first confirmed this observation in our settings by determining the minimal inhibitory concentration of antibiotics on planktonic bacteria (minimal inhibitory concentration, MIC) and on biofilm formation (biofilm prevention concentration, BPC_90_). For *A. baumannii*, we measured a 128 to over 512 times higher BPC_90_ than the corresponding MIC for all tested antibiotics (**Table 1**). The effect was less striking for *P. aeruginosa* with a BPC_90_/MIC ratio of 2-4 (**Table 1**).

**Table 1.**

| Antibiotic/AMP | MIC (µg/ml) | BPC<sub>90</sub> (µg/ml) | MBIC (µg/ml)<br>Resazurin / CV | MBEC<sub>90</sub> (µg/ml) |
| --- | --- | --- | --- | --- |
| *A. baumannii* ATCC 19606 |  |  |  |  |
| Ciprofloxacin | 1 | > 256 | > 256 | > 256 |
| Tetracycline | 1 | 128-256 | > 256 | > 256 |
| Gentamicin | 32 | > 256 | > 256 | > 256 |
| Levofloxacin | 0.5 | > 256 | > 256 | > 256 |
| TAT-RasGAP<sub>317-326</sub> | 8 | 8-16 | 32 | > 256 |
| Polymyxin B | 4 | 8-16 | 32 / 128 | > 256 |
| Melittin | 16 | 16 | 128 | > 256 |
| *P. aeruginosa* PA14 |  |  |  |  |
| Ciprofloxacin | 0.2 | 0.4 | < 2 / > 256 | > 256 |
| Tetracycline | 16 | 8-32 | 32-64 / > 256 | > 256 |
| Gentamicin | 4 | 8-16 | 32 / > 256 | > 256 |
| TAT-RasGAP<sub>317-326</sub> | 32 | 32 | 64 | > 256 |
| Polymyxin B | 2 | 2-4 | 32 | > 256 |
| Melittin | 64 | 128 | > 256 | > 256 |
| *S. aureus* ATCC 29213 |  |  |  |  |
| Ciprofloxacin | 0.4 | 128-256 | > 256 | > 256 |
| TAT-RasGAP<sub>317-326</sub> | 128 | 128-256 | > 256 | > 256 |

MIC: minimal inhibitory concentration, BPC: biofilm prevention concentration, MBIC: minimal biofilm inhibitory concentration, MBEC: minimal biofilm eradication concentration, CV: crystal violet. For MBIC, the indication of a single value means that the result was the same with resazurin and CV assays.

In a further step, we tested whether these antibiotics could limit biofilm expansion (measuring the minimal biofilm inhibitory concentration, MBIC) and eradicate established biofilms (measuring the minimal biofilm eradication concentration, MBEC_90_). We used the reduction of resazurin to resofurin as a surrogate of bacterial viability and measured total biomass with crystal violet staining. All the tested antibiotics were ineffective at both inhibiting expansion and eradicating *A. baumannii* biofilms (MBIC and MBEC_90_ > 256 µg/ml, **Table 1**). While ciprofloxacin, tetracycline and gentamicin inhibited the proliferation of viable bacteria in *P. aeruginosa* biofilms at concentrations corresponding to 2-4 times the BPC_90_, we observed no effect on biomass (**Table 1**). These results confirm the low efficiency of classical antibiotics on biofilms.

### TAT-RasGAP_317-326_ potently inhibits the formation and expansion of *A. baumannii* and *P. aeruginosa* biofilms

Well-described AMPs such as melittin and polymyxin B efficiently reduce bacterial biofilm formation and the biomass of established biofilms [27, 28]. In line with published results, we observed a consistent inhibitory effect of these AMPs on *A. baumannii* and *P. aeruginosa* biofilm formation. The BPC_90_ of melittin was equal to the MIC for *A. baumannii* biofilms and 2-4 times higher than the MIC for *P. aeruginosa* biofilms (**Table 1**). For polymyxin B, the BPC_90_/MIC ratios were 2-4 and 1-2 for *A. baumannii* and *P. aeruginosa* respectively (**Table 1**). As TAT-RasGAP_317-326_ is efficient against planktonic *A. baumannii* and *P. aeruginosa*, we hypothesized that it would also potently inhibit the formation of biofilms. TAT-RasGAP_317-326_ inhibited *A. baumannii* biofilm formation at 8-16 µg/ml corresponding to 1-2 times the MIC (**Fig. 1A**-**B** and **Table 1**). The BPC_90_ of TAT-RasGAP_317-326_ on *P. aeruginosa* biofilms was equal to its MIC (32 µg/ml; **Fig. 2A-B** and **Table 1**).

**Figure 1:**
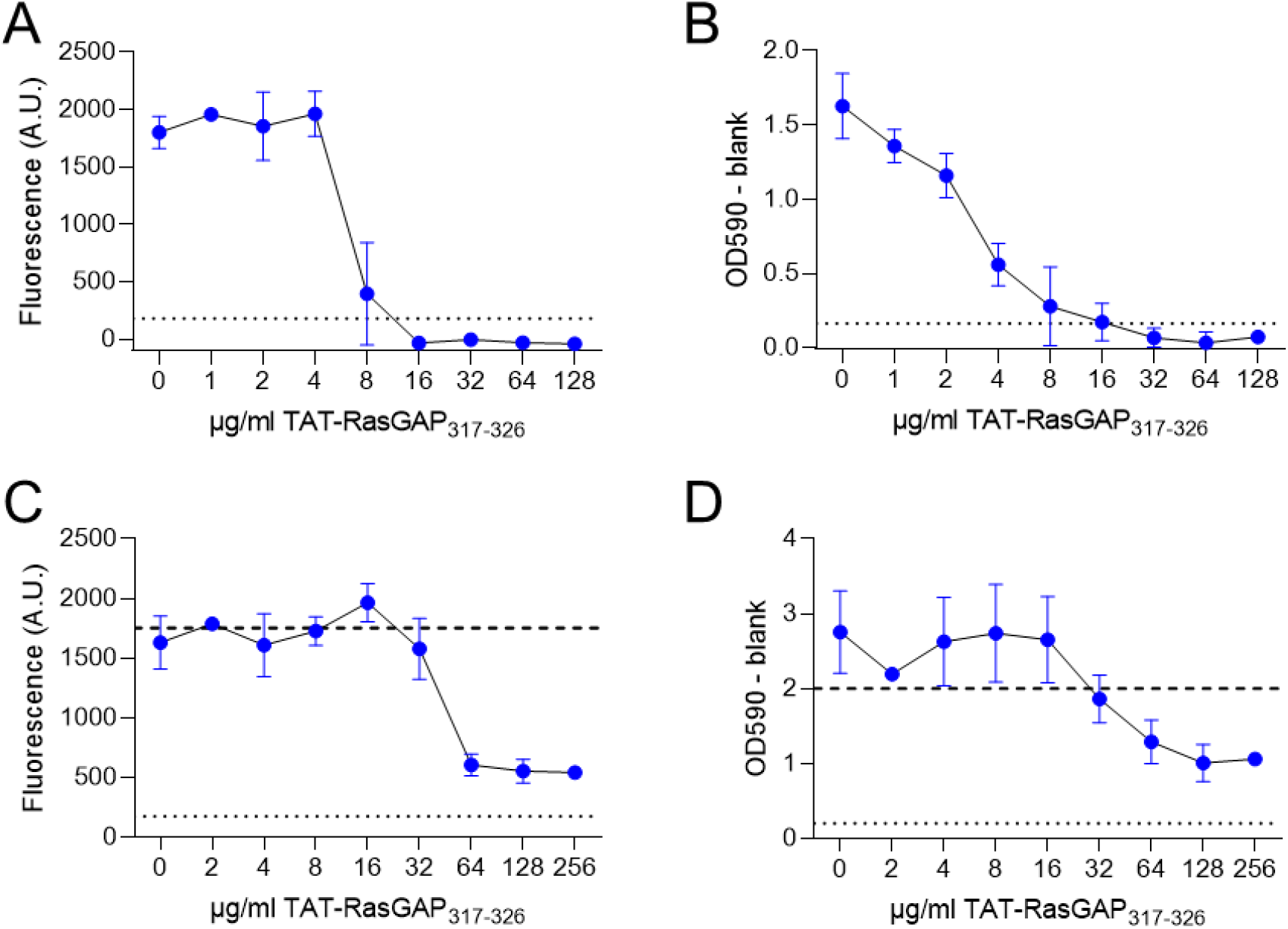
TAT-RasGAP_317-326_ inhibits *A. baumannii* biofilm formation and expansion. The effect of TAT-RasGAP_317-326_ on biofilm formation (A,B) and on mature biofilm expansion (C,D) was assessed with resazurin reduction (A,C) and crystal violet (B,D) assays. Dotted lines indicate 10% of initial signal. Dashed lines indicate values for mature biofilms before treatment. Data are means ± SD of three experiments performed in quadruplicate.

**Figure 2:**
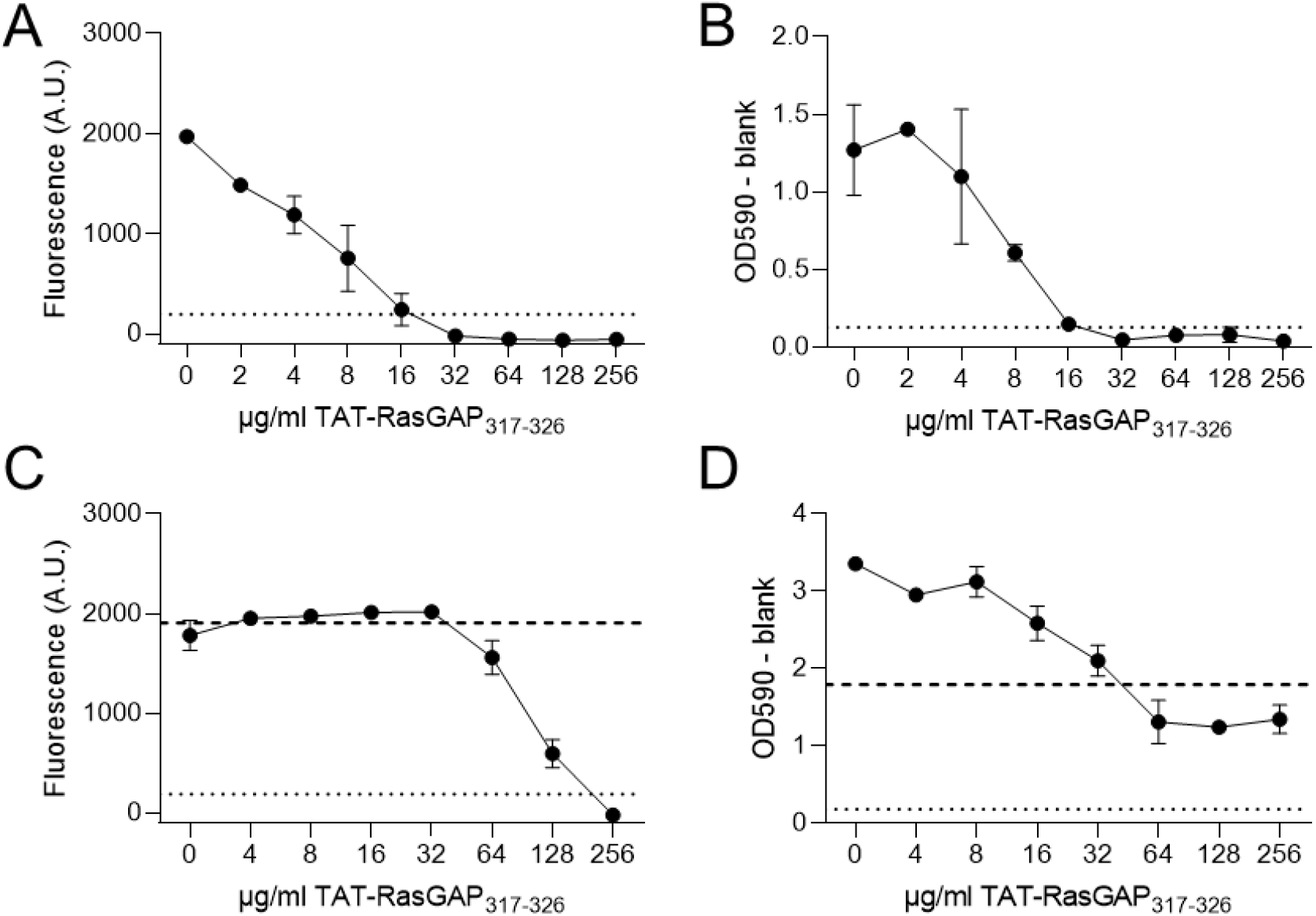
TAT-RasGAP_317-326_ inhibits *P. aeruginosa* biofilms formation and expansion. TAT-RasGAP_317-326_ was added simultaneously with bacteria (A-B) to measure inhibition of biofilm formation, or on mature biofilm (C-D) to assess biofilm expansion limitation. Bacterial viability was approximated with resazurin reduction assay (A,C) and biofilm biomass with crystal violet staining (B,D). Dotted lines indicate 10% of initial signal. Dashed lines indicate values for mature biofilms before treatment. Data are means ± SD of three experiments performed in quadruplicate.

**Figure 3:**
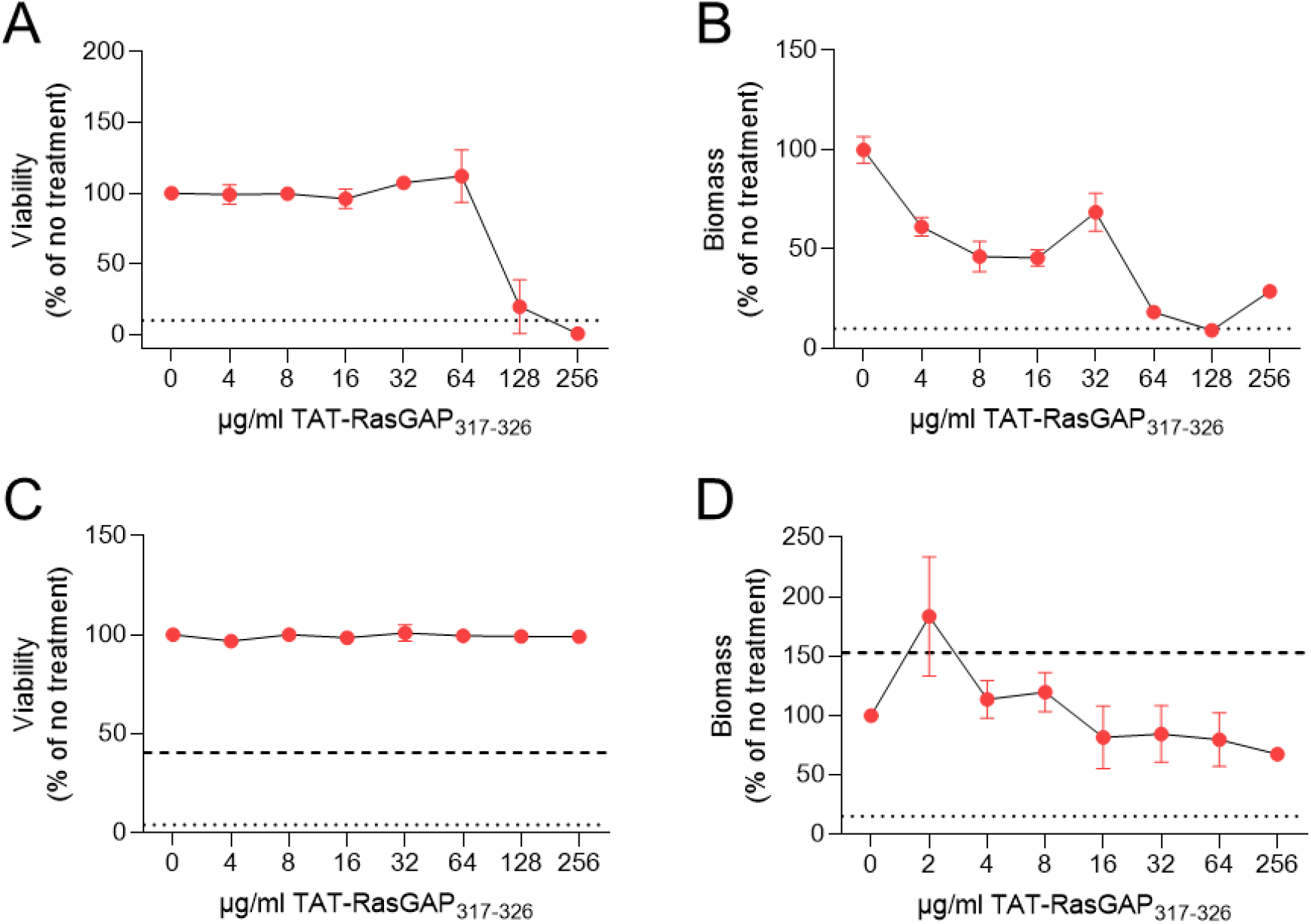
TAT-RasGAP_317-326_ reduces *S. aureus* biofilm formation. (A-B) Addition of TAT-RasGAP_317-326_ at times of biofilm induction reduced the viability (A) and the biomass (B) of *S. aureus*. (C-D) Treatment of mature biofilms with TAT-RasGAP_317-326_ had no effect on bacterial viability (C) and biomass (D). Viability was measured with resazurin (A,C) and biomass with crystal violet (B,D) assays. Dotted lines indicate 10% of initial signal. Dashed lines indicate values for mature biofilms before treatment. Data are means ± SD of three experiments performed in quadruplicate.

Since classical antibiotics had little to no effect on biofilm expansion and eradication (**Table 1**), we tested the effect of AMPs on established biofilms. TAT-RasGAP_317-326_ limited the expansion of *A. baumannii* biofilms with a MBIC/MIC ratio of 4 (**Fig. 1C-D** and **Table 1**). Melittin was effective at 8 times the MIC, while Polymyxin B had a MBIC/MIC ratio of 8 for viability and 32 for biomass (**Table 1**). TAT-RasGAP_317-326_ efficiently inhibited *P. aeruginosa* biofilm expansion with a MBIC/MIC ratio of 2 (**Fig. 2C-D** and **Table 1**). In contrast, polymyxin B had a higher MBIC/MIC ratio of 16 while the MBIC of melittin could not be determined, being higher than 256 µg/ml (**Table 1**). However, none of the tested AMPs had the potency to eradicate more than 90% of *A. baumannii* or *P. aeruginosa* biofilms at the maximal tested concentration of 256 µg/ml.

### TAT-RasGAP_317-326_ inhibits the formation of *S. aureus* biofilms

TAT-RasGAP_317-326_ has an effect on both Gram-negative and Gram-positive bacteria in planktonic cultures [20]. We thus tested its impact on *S. aureus* biofilms. Despite the high MIC of TAT-RasGAP_317-326_ on planktonic *S. aureus* (128 µg/ml), we could measure a BPC_90_/MIC ratio of 2 (**Table 1**). In comparison, the BPC_90_/MIC ratio of ciprofloxacin was 320-640 (**Table 1**). However, as expected due to the high MIC and BPC_90_ of TAT-RasGAP_317-326_, the MBIC and MBEC_90_ on *S. aureus* biofilm could not be estimated being higher than 256 µg/ml.

## Discussion

Bacterial biofilms cause nosocomial infections and underlie several chronic infections. The complex structure of biofilms renders them refractory to treatments with classical antibiotics. Here we show that the AMP TAT-RasGAP_317-326_ has a potent inhibitory effect on *A. baumannii, P. aeruginosa* and, to a lesser extent, *S. aureus* biofilm formation and expansion. These observations support the potential of AMPs as alternative to classical antibiotics in anti-biofilm treatment.

In order to test the quality of the biofilm we produced *in vitro*, we measured its resistance to classical antibiotics. As reported in literature, we observed increased resistance of *A. baumannii* and *P. aeruginosa* biofilms to classical antibiotics. *A. baumannii* biofilms formed in the presence of concentrations of antibiotics equivalent to 128 to more than 512 times the MIC on planktonic bacteria (**Table 1**), while *P. aeruginosa* biofilm formation was abolished with concentrations equivalent to 2-4 times the MIC. This difference in biofilm resistance to antibiotics is possibly caused by a faster resistance development during *A. baumannii* biofilm formation, partly triggered by the antibiotics themselves [29]. While antibiotics showed some effect on biofilm formation at moderate to high concentrations, they were not efficient in inhibiting proliferation or disrupting mature biofilms (**Table 1**, [30]). This may be due to the low permeability of the extracellular matrix to classical antibiotics, and highlights the need of alternative strategies to treat infections involving biofilms.

In contrast to antibiotics, TAT-RasGAP_317-326_, polymyxin B and melittin limited both *A. baumannii and P. aeruginosa* biofilm formation and expansion. Similar promising results have been observed with other AMPs including the human AMP LL-37 and the bacterial bacteriocins [31, 32]. Moreover and despite its high MIC, TAT-RasGAP_317-326_ also reduced *S. aureus* biofilm formation. These results are encouraging and suggest that combinations of TAT-RasGAP_317-326_ with other drugs might be helpful in the treatment of biofilm-associated infections. As biofilm are composed of heterogeneous bacterial subpopulations, the combination of AMPs with antibiotics is a valuable way to avoid the survival of persister cells. The combination of melittin and LL-37 with classical antibiotics reduced the corresponding MBEC several folds [33]. We need to test in future studies the anti-biofilm effect of combinations of TAT-RasGAP_317-326_ with classical antibiotics. Indeed, TAT-RasGAP_317-326_ might have advantages compared to other AMPs: its quite low toxicity to mammalian cells [20] and its chimeric nature against which bacteria should not have developed resistance. Moreover, since biofilms are often composed of multiple bacterial species, the broad-spectrum antimicrobial activity of TAT-RasGAP_317-326_ is interesting for future clinical developments. The current limitation of TAT-RasGAP_317-326_ for clinical use is its high excretion rate and subsequent low bioavailability [20]. We thus need to develop strategies to increase the distribution of this peptide *in vivo*, which could be achieved by chemical modifications or specific delivery methods.

In summary, we show in this report that the AMP TAT-RasGAP_317-326_ has potent anti-biofilm activity *in vitro* against *A. baumannii* and *P. aeruginosa* biofilms and is relatively active on *S. aureus* biofilms. This makes TAT-RasGAP_317-326_ a promising tool in the treatment of biofilm-associated infections and might lead to the development of new strategies to prevent biofilm formation.

## Authorship

TH and SH performed experiments. TH, MG, CW and NJ were involved in the planning of the project and discussed the results. TH analysed the results. TH and NJ wrote the manuscript.

## Acknowledgements

NJ and CW received an interdisciplinary grant from the University of Lausanne. We thank Prof. Gilbert Greub for providing laboratory equipment. We thank the Prof. Nibbering group (Leiden, Netherlands) for their assistance in the implementation of the protocol of *in vitro* formation of biofilms and Prof. Eberl (Zurich, Switzerland) for providing strains.

